# Male and Female Brain Coherence Models of Cognitive Performance and Psychopathology

**DOI:** 10.1101/2022.09.28.509939

**Authors:** Magda L. Dumitru, Max Korbmacher, Hauke Bartsch

**Author notes:** Corresponding Author: Magda L. Dumitru, PhD, Department of Biological and Medical Psychology, Jonas Lies vei 91, 7807, Bergen, Norway.

## Abstract

Finding reliable imaging biomarkers of mental illness has been a major challenge, on a par with the quest for biomarkers of the male versus the female brain, as the two types of imaging inform one another. We explored the hypothesis that the degree of coherence (internal isomorphism) between brain volumes of the left versus the right hemisphere for patients with psychopathological conditions follows the brain coherence pattern of the healthy male or healthy female. We developed the ‘distance index’ (DI) as a biomarker of brain coherence and compared it with three ad hoc coherence measures. We found that only DI could reliably distinguish males from females and patients from controls. Also, cortical regions with highest DI scores were swapped between males and females across groups following male/female models of psychopathology. Furthermore, although indices were similar in predicting cognitive performance, DI provided a more proportionate prediction pattern across diagnosis groups, and more robust interactions with males/females. These findings highlight the importance of brain coherence, particularly measured by DI, for phenotyping sex and mental illness.

---

The last few decades have seen a relentless quest for biomarkers of psychopathological disorders using magnetic resonance imaging (MRI). Indeed, finding reliable indicators of mental illness could help clarify underlying physiological mechanisms as well as associated risk and protective factors, which may further improve treatment and prevention. Nevertheless, the results have been modest for reported characteristics of cortical and subcortical brain volumes in patients with psychopathologies (Lee et al., 2013; Cross-Disorder Group of the Psychiatric Genomics Consortium, 2013). At the same time, research on differences in brain structure between males and females in the context of psychopathology have not gone beyond the classic dichotomy between the less asymmetric female brain and psychopathological brain on the one hand, and the more asymmetric male brain on the other hand. However, these findings draw on conflicting MRI evidence. So, for example, sexually dimorphic brain asymmetries were reported for cortical regions subserving language, in particular the ‘planum temporale’ (e.g., de Courten-Myers, 1999; Good et al., 2001; Shapleske et al., 1999), but whereas some of them suggest stronger leftward lateralisation in males (Guadalupe et al., 2015), others imply leftward lateralisation in females (Ruigrok et al. 2014). After controlling for total intracranial volume, few if any statistically significant differences in grey matter volumes are left standing between the sexes (cf. review by Sanchis-Segura et al., 2019), despite statistically higher brain volumes in males compared to females. Renewed interest in finding successful measures of male/female structural differences may explain the onset and timeline of psychopathological conditions as well as of specific cognitive abilities, which differ between the sexes (Halpern, 2012). So, for example, Attention Deficit and Hyperactivity Disorder (ADHD) is more common in males, whereas depression is more common in females (Angold, Costello, and Worthman, 1998).

## Brain asymmetry measures

Brain asymmetry has been described centuries ago (Eberstaller, 1889; for a review see Toga and Thompson, 2003) in connection with the significant expansion of the human neocortex, which gradually saw the emergence of evolutionary advantages such as enhanced information processing (Dadda and Bisazza, 2006) and high-level cognitive functions. Indeed, the two halves of the brain are nowadays slightly asymmetric, with the right side warped somewhat forward relative to the left side, as part of the so-called “Yakovlevian torque”. This rotation has favoured lateralized structure and functionality, for example the development of a longer superior boundary of the left temporal lobe to afford language abilities. Overall, the asymmetry between the left and the right side of the brain, which is well-documented in various species, is more pronounced in humans compared to other primates mainly due to the expansion of specialized cortical areas (Wang, Buckner, & Liu, 2014). In contrast, similarities across the two brain hemispheres have been associated with psychopathological conditions such as schizophrenia, ADHD, or bipolar disorder (Crow et al. 1989; Geschwind & Levitsky, 1968; Hajima et al. 2013; Postema et al. 2021; Sommer et al., 2008).

Anatomical differences between the left and the right hemisphere are routinely calculated using the “laterality index” (LI), which involves pairwise items and is given by the formula (left - right)/ (left + right). LI helps determine, for example, whether the left temporal lobe is larger than the right, or whether the right occipital lobe is larger than the left, without considering the impact of other (related) brain areas. In other words, LI lacks sensitivity for the ‘larger picture’, which is nevertheless specific to psychopathological conditions, where more than one region is usually responsible for specific symptoms. Several studies using LI have found differences in the Yakovlevian torque for depression, bipolar disorder, and schizophrenia (Mock et al., 2012; Maller et al., 2014; Maller et al., 2015), but results are inconclusive, possibly because of additional variation that remains unaccounted for. Identifying a biomarker sensitive to volumetric differences between a specific region of interest and multiple regions of interest within a structure is more likely to increase precision because it covers a larger volume and is therefore less noisy, in addition to being more ecologically valid (mental illness usually involves several brain areas). At the other end of the analysis spectrum, even minor departures from normal brain asymmetries have sometimes been associated with mental disorders (Petty, 1999), though defining normal individual variability and underlying mechanisms remains challenging. Again, limited-precision analyses may be responsible for these outcomes, which explains why progress in MRI-driven search for biomarkers of sex and psychopathological conditions has been rather slow.

## The distance index DI

We set out to investigate a novel imaging biomarker to show that, when properly measured, brain coherence can reliably predict both sex and psychopathology. In the broadest sense, coherence can be understood as internal isomorphism of whole-brain volumes, which can be captured by various coherence measures, as detailed further below. Our specific hypothesis is that the degree of coherence between brain volumes of the left versus the right hemisphere for patients with psychopathological conditions follows the brain coherence pattern of the healthy male or healthy female. We hypothesized that brain structure may vary at several sites in the left and the right hemispheres across sexes and diagnosis groups, and these variations depend on each other, thus creating patterns of asymmetries that are left unexplained by current analysis methods.

We explored imaging biomarkers associated with brain coherence that capture these patterns. To measure coherence, we computed a ‘distance index’ (DI) over grey matter volumes for each region of interest and contrasted it with three other coherence measures using the Desikan-Tourville atlas (Desikan et al. 2006). Unlike lateralization measures reported in the literature, DI weighs the volume of each substructure relative to the volumes of all other substructures, as described in Figure 1 below. Here, DI spans pairs of vectors over the 31 regions in each brain hemisphere, thus deriving differences between all regions instead of pairwise weightings, as was routinely the case when calculating differences in symmetry (i.e., when using LI).

**Figure 1.**
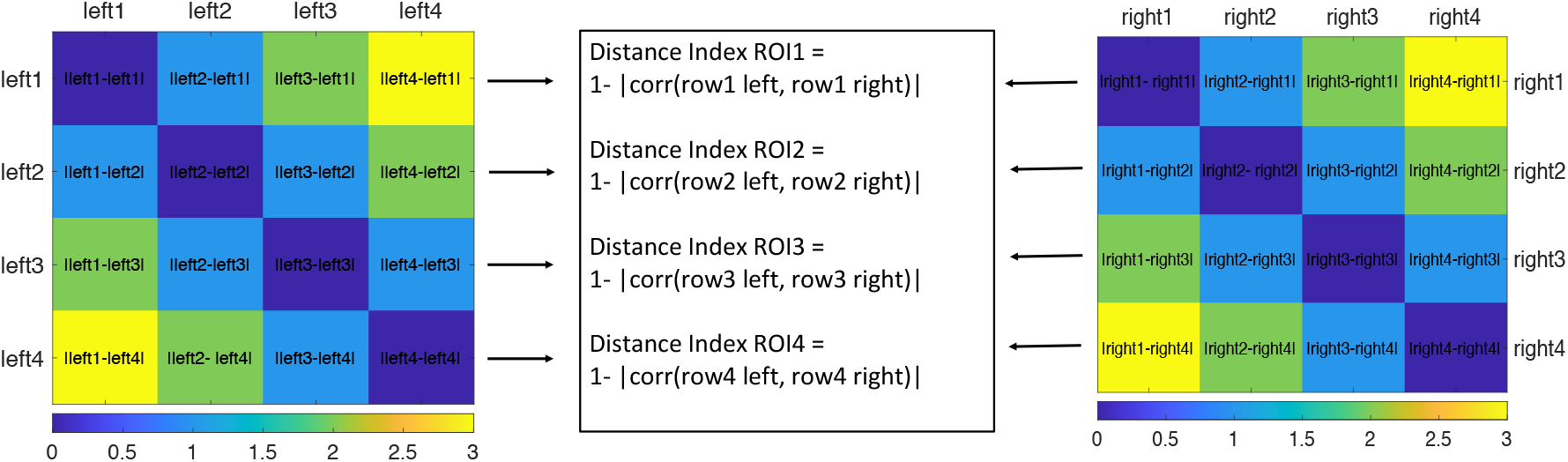
Calculating the distance index DI. First, we obtained a matrix of absolute differences between each ROI volume and the remaining 30 ROI volumes in the left hemisphere, and then a second matrix calculated in the same way for ROIs in the right hemisphere. Here, the matrices have equal values. Next, we obtained Pearson correlation coefficients between the two matrices and subtracted each coefficient from 1, to obtain a vector of differences. DI in the form of absolute differences were derived for each difference, representing overall magnitudes. The higher the DI, the lower the coherence between the left and right region of interest.

We compared ROI scores for males versus females for each of the four coherence indices and compared groups (ADHD, Bipolar, Schizophrenia, and Controls) in ANOVAs for each index and predicted significant differences between sexes that vary with diagnosis. For example, coherence could be lower in males than in females for healthy controls, but higher in females than in males for individuals with ADHD, since ADHD is more prevalent in males than in females (Novik et al., 2006; Scahill & Schwab-Stone, 2000), and therefore we expect it to follow a reversed (male-specific) brain structure pattern. Second, identifying brain regions with low coherence in males with ADHD and in females from the control group, for example, would provide support for our hypothesis that male- and female-typical brain structures serve to predict psychopathology. Third, by projecting coherence scores on the cortex, we could identify a shared coherence network. Finally, we performed principal component analysis (pca) and entered the four components with highest eigenvalues for each index as predictors in linear models of cognitive scores in 7 questionnaires. We predicted that, if brain coherence were relevant to cognitive performance, we would observe significant interactions between coherence indices and diagnosis groups and/or sex, which would further indicate that the brain structure specific of psychopathological conditions simulates the structure of the male brain or the female brain. In other words, the degree of brain coherence specific to males, for example, could also be specific to individuals suffering from certain psychopathologies, whereas the degree of brain coherence specific to females could also be specific to individuals suffering from other psychopathologies.

## Methods

### Participants

We analyzed MRI and behavioural data collected as part of the neuroimaging project by the UCLA Consortium for Neuropsychiatric Phenomics included 40 individuals with ADHD (20 females), 49 individuals with bipolar disorder (21 females), 44 individuals with schizophrenia (10 females), and 58 healthy individuals (26 females), all right-handed. Additional participant information can be found on the project webpage (Gorgolewski et al. 2017; Poldrack et al. 2016).

### MRI data acquisition

Neuroimaging data were acquired on a 3T Siemens Trio scanner. T1-weighted high-resolution anatomical scans (MPRAGE) were collected with the following parameters: slice thickness = 1mm, 176 slices, TR=1.9s, TE=2.26ms, matrix=256 × 256, FOV=250mm. Cortical reconstruction, volumetric segmentation, and parcellation using the Desikan-Killiany atlas (Desikan et al. 2006) were performed with the *Freesurfer 6* image analysis suite, which is documented and freely available for download online (http://surfer.nmr.mgh.harvard.edu/). Standard cross-sectional processing stream was used, which included motion correction and averaging, automated Talairach transformation, segmentation of the subcortical white matter and gray matter volumetric structures, intensity normalization, tessellation of the gray matter white matter boundary, automated topology correction, and surface deformation following intensity gradients.

### Data analysis

#### Calculating coherence indices DI, LI, SI, and ZI

Based on average brain volumes following the Desikan-Tourville atlas (Desikan et al. 2006), we derived four coherence measures. First, we calculated the novel DI measure as detailed in Figure 1. We started by deriving a series of absolute differences between the volume of each region of interest and the volumes of the remaining 30 regions of interest for the left hemisphere. We then applied the same calculations to the right hemisphere, thus obtaining two 31 × 31 matrices, one for the left, and the other for the right hemisphere, as seen in Figure 1. We correlated each vector in the two matrices and its correspondent with each other, then subtracted each Pearson correlation coefficient from 1 for each region and for each hemisphere. A higher DI for a particular region would indicate greater differences between the left and the right hemisphere, hence lower coherence, whereas a lower DI would indicate fewer differences, hence higher coherence between hemispheres.

Next, we calculated a second coherence measure, namely a modified version of the laterality index LI, by subtracting the volume of each region of interest in the left hemisphere from its counterpart volume in the right hemisphere, before dividing this number by the sum of the two volumes, according to the classic formula (L-R)/(L+R). In a last step, we obtained absolute values, as we aimed to measure the magnitude of the left-right difference instead of identifying which region, to the left or to the right, has a greater volume than the other, as per usual. The third measure of coherence we considered was a straightforward subtraction index SI, which we obtained by subtracting the volume of one region of interest from the other, before deriving the absolute value of this difference. As with LI, we aimed to measure the magnitude of the left-right difference when calculating SI, instead of ranking the magnitudes of the two regions. The fourth and last measure of coherence we included was the z-scored laterality index ZI, which provides z-score values for measures calculated using LI. In sum, the four indices provide coherence estimates for the same volumes – in this case between homologous regions of interests in the left and the right hemispheres in the form of positive differences in magnitude. In the case of DI, these differences involve all regions of interest for each volume investigated, whereas in the case of LI, SI, and ZI, these differences are established between pairs of homologous left-right volumes. The Matlab code we used for calculating the four indices is available on GitHub (https://github.com/magda47/Distance-Index).

We obtained, for each index, a score for each region of interest, hence 31 scores for all ROIs, separately for males and for females, resulting in two sets of scores for each diagnosis group. By comparing them in statistical analyses, we aimed to identify the most robust biomarker of sex and psychopathology. Specifically, we aimed to determine whether the underlying coherence mechanism is recurrent involvement of all brain regions in estimating each brain region, as with DI, or rather ranked pairwise relations between brain regions, as with LI, or yet again the ranking of single brain regions and their counterparts, as with SI or, finally, average-corrected estimations of region pairs, as with ZI.

#### Cortical maps of DI, LI, SI, and ZI as coherence measures

To determine the best biomarker of the male or female brain and corresponding psychopathological conditions, we entered the four indices into *t*-tests over male and female scores for each group, and in Friedman’s two-way ANOVA by ranks over diagnosis groups. Next, we constructed brain maps for the top 20% highest index values in males and females for each group (ADHD, Bipolar, Schizophrenia, and Controls). Since ADHD is far more prevalent in males than in females (Novik et al., 2006; Scahill & Schwab-Stone, 2000), finding brain regions that are swapped between males and females depending on diagnosis group that is, specific of males in ADHD but specific of females in the other groups and the other way around would provide support for our hypothesis that brain structures typical to males or to females predict psychopathology. Coherence maps for regions with highest scores for each index may further overlap, highlighting method-independent regions of coherence between the hemispheres.

#### Coherence-based predictors of cognitive test scores

To identify which of the four indices and associated coherence map is the closest predictor of sex and diagnosis (ADHD, BP, SZ, and controls), we entered the highest four components into linear models as predictors of cognitive test scores alongside sex, diagnosis group, and their interaction with each component. We compiled scores from 25 measures of 7 cognitive tests: total adjusted pumps on the Balloon Analog Risk Task (BART); total correct Go responses, false alarms, and mean RT correct Go responses on the Continuous Performance Test (CPT); list B free recall, free recall intrusions, recognition discriminability, short delay cued recall, total intrusions, long delay free recall, long delay cued recall, short delay free recall, learning slope, and long delay free recall on the California Verbal Learning Test-II (CVLT); total errors and total correct responses on the Delis-Kaplan Executive Function System (D-KEFS); letter-number sequencing, matrix reasoning, and vocabulary on the Wechsler Adult Intelligence Scale (WAIS); accuracy and reaction time for responses on the Stroop Color-Word Task (STROOP); mean reaction time on correct GO trials, percent inhibition, percent GO response, and direction errors on the Stop Signal Task (SST). In sum, the seven cognitive tests measured language, memory, attention, cognitive control, risk-taking, and general intelligence. Core measures associated with these tests overlap in their potential to estimate specific cognitive abilities, as we aimed to increase the chances of finding significant predictors among indices and their interaction with sex and diagnosis group.

## Results

### Direct comparison of DI, LI, SI, and ZI coherence measures

Figure 2 summarizes average scores across coherence indices DI, LI, SI, and ZI. We calculated paired *t*-tests over the 31 regions of interest between males and females for each index and for each diagnosis group (ADHD, Bipolar, Schizophrenia, and Controls); results are summarized for each index. DI differences between the left and right hemisphere were significantly greater for males compared to females, suggesting that the brain of females is more coherent than the brain of males, for the Bipolar, Schizophrenia, and Control groups. For the ADHD group, the results were reversed, such that DI scores were higher for females than for males suggesting that, depending on diagnosis group, brain coherence can be higher in males compared to females, or in females compared to males. For the other indices, differences between males and females were not significant, with a few exceptions showing higher coherence in females compared to males, as for DI: in the Schizophrenia group for LI, and in the Bipolar and Schizophrenia groups for SI.

**Figure 2.**
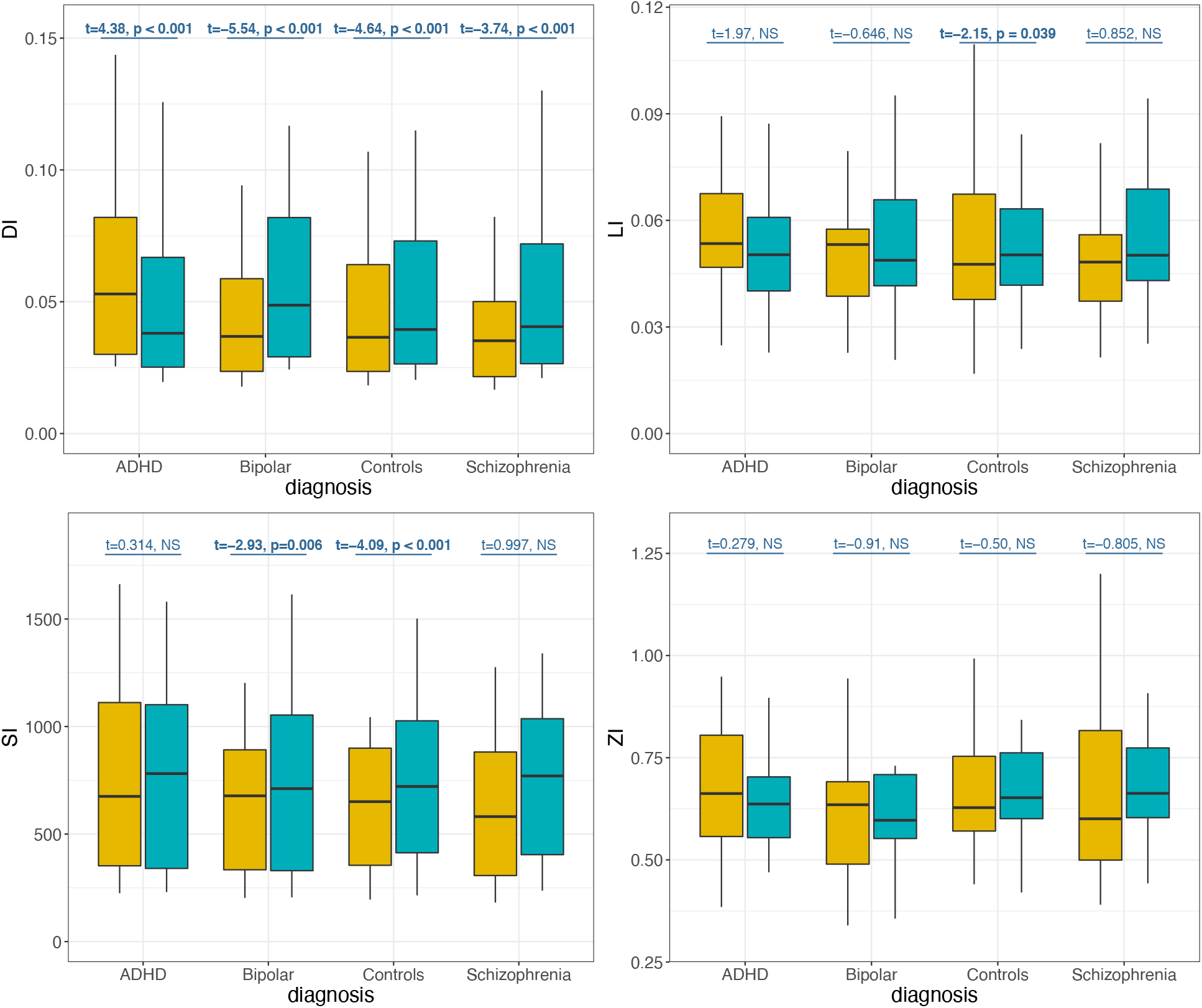
Brain coherence values across males and females. Paired *t*-tests values are given for each coherence index (DI, LI, SI, and ZI) over all groups (ADHD, Bipolar, Schizophrenia, and Controls). Boxplots are gold yellow for females, and blue for males.

We further conducted related-samples Friedman’s two-way ANOVA by ranks to determine whether coherence differs significantly among groups (ADHD, BP, SZ, and Controls) for each index. We found it to be the case for DI, *χ2*(3) = 47.013, *p* < 0.001. Bonferroni-adjusted pairwise comparisons revealed higher coherence, hence lower DI values for controls compared to BP (*M* = 0.0479 vs. 0.052, *p* < 0.001) and to ADHD (*M* = 0.0479 vs. 0.0552, *p* < 0.001). The remaining comparisons were (close-to) significant but did not survive adjustment. Overall, controls had the lowest DI. For the other indices, results were not significant: for LI, *χ2*(3) = 4.742, *p* = 0.192. For SI, *χ2*(3) = 5.284, *p* = 0.152. For ZI, *χ2*(3) = 3.619, *p* = 0.306.

### Cortical maps of DI, LI, SI, and ZI coherence measures

We constructed coherence maps for the 20% highest-scoring brain regions across males and females, as seen in Figure 3. For each index, we identified regions that were common across males and females for each index, as follows. For DI, the ROIs common between sexes for ADHD were the supramarginal, inferior parietal, and postcentral gyrus; the rostral middle frontal was specific to males, and the superior parietal was specific to females. For DI Bipolar, the supramarginal, inferior parietal, postcentral, and fusiform were common for males and females, with the rostral middle frontal specific to females and the superior parietal specific to males. We can already identify the latter two areas as being swapped between sexes with respect to ADHD areas. Further, the same areas common to males and females in the Bipolar group were also common to males and females in the Schizophrenia group. In addition, we found the rostral middle frontal specific to females, as in the Bipolar group, and the inferior temporal specific to males. For the DI Controls group, the supramarginal, inferior parietal, and postcentral area were common across males and females, with the rostral middle frontal and the caudal middle frontal specific to females, and the fusiform and the lingual specific to males. Overall, DI-defined ROIs were similar across the sexes, with a double swap of the superior parietal and the rostral middle frontal across the ADHD and Bipolar group. The latter males/females swap further applied between ADHD and all remaining diagnosis groups.

**Figure 3.**
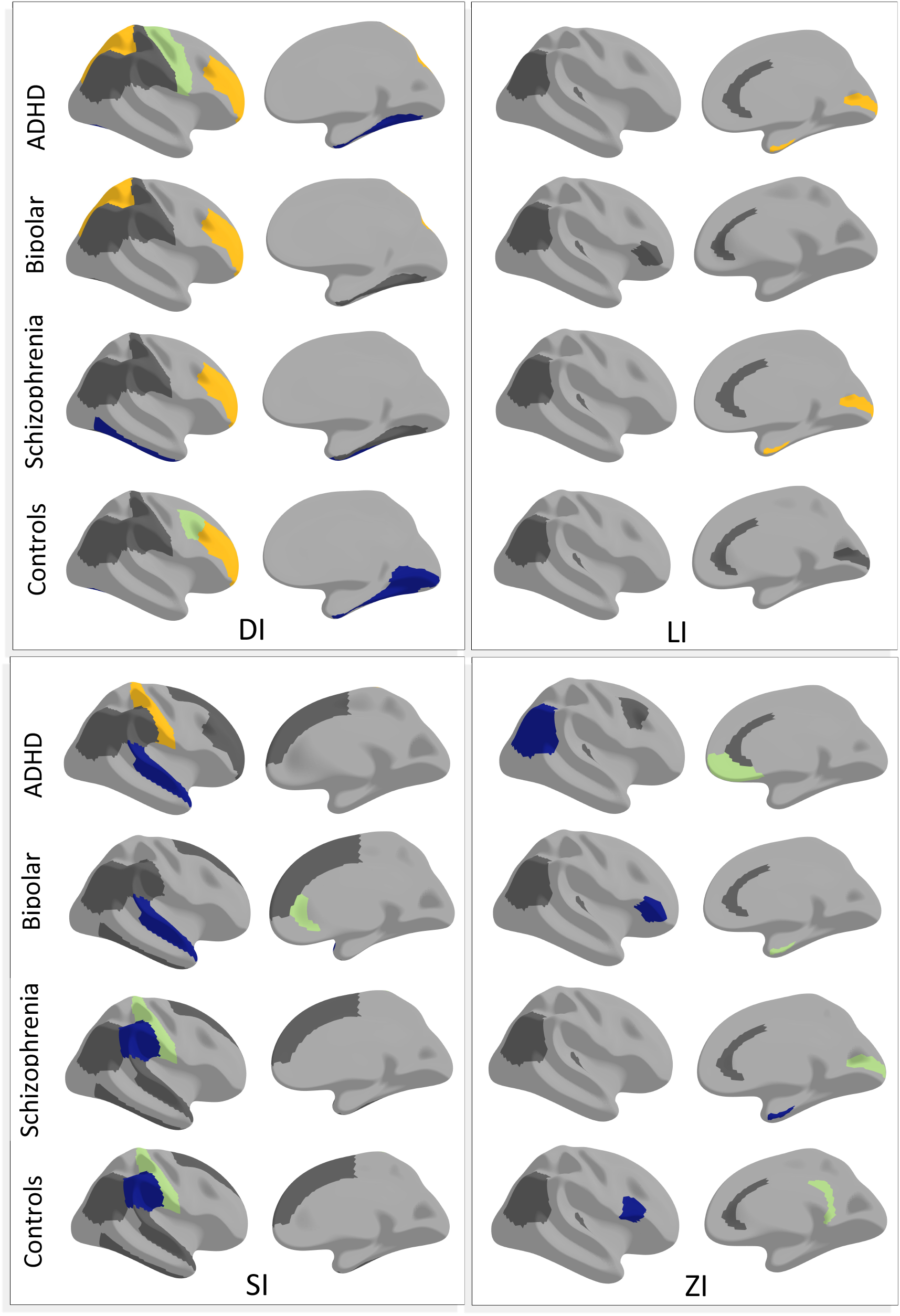
Coherence maps for males and females across groups and indices. ROIs common between males and females in dark grey; ROIs switched between ADHD and other groups in yellow; ROIs specific to females in olive green; ROIs specific to males in navy.

For the LI ADHD group, ROIs were common between sexes for the rostral anterior cingulate, transverse temporal, caudal anterior cingulate, and inferior parietal, with the entorhinal specific to females, and the pericalcarine specific to males. For the LI Bipolar group, all ROIs were common across sexes, and included the rostral anterior cingulate, caudal anterior cingulate, transverse temporal, and inferior parietal. For the LI Schizophrenia group, common areas were the rostral anterior cingulate, caudal anterior cingulate, transverse temporal, and inferior parietal. Further, the pericalcarine was specific to females, whereas the entorhinal was specific to males, in a double swap with respect to the ADHD group. For the LI Controls group, all ROIs were common across sexes, and included the rostral anterior cingulate, caudal anterior cingulate, transverse temporal, pericalcarine, and inferior parietal. Overall, LI-based areas were common between males and females, with a double region swap between sexes in the ADHD and Schizophrenia group.

For the SI ADHD group, four ROIs were common between sexes: the superior frontal, rostral middle frontal, supramarginal, and inferior parietal. The postcentral was specific to females, and the superior temporal was specific to males. For the SI Bipolar group, we found again four common areas between sexes: the superior frontal, supramarginal, inferior temporal, and inferior parietal, with the rostral anterior cingulate specific to females, and the superior temporal specific to males. For the SI Schizophrenia group, there were again four common areas between sexes, the superior frontal, inferior temporal, superior temporal, and inferior parietal, with the postcentral specific to females, and the supramarginal specific to males. For the SI Controls group, common areas between sexes were the superior frontal, superior temporal, and inferior parietal. There were two regions specific to females, the supramarginal and the lateral occipital, and two regions specific to males, the rostral middle frontal and the postcentral. The latter was a swap relative to females in the ADHD group. Overall, SI-based areas were common between males and females, with a double region swap between sexes in the ADHD and Controls group.

For the ZI ADHD group, there were four ROIs common across sexes: the rostral anterior cingulate, transverse temporal, caudal anterior cingulate, and caudal middle frontal. The medial orbitofrontal was specific to females, and the inferior parietal was specific to males. For the ZI Bipolar group, there were again four areas common between males and females: the rostral anterior cingulate, caudal anterior cingulate, transverse temporal, and inferior parietal. The entorhinal was specific to females, and the pars triangularis was specific to males. For the ZI Schizophrenia group, the same four regions were common across sexes as in the Bipolar group. In addition, the pericalcarine was specific of females, whereas the entorhinal was specific of males. For the ZI Controls group, the same four areas were common between males and females as in the Bipolar and the Schizophrenia group. In addition, the isthmus cingulate was specific to females, and the pars opercularis was specific to males. Overall, ZI-based areas were largely common between males and females without any swaps between males and females across groups. Among the ROIs common between males and females across groups and across indices, we singled out the inferior parietal as a key region for measuring brain coherence.

### Coherence-based predictors of cognitive scores

We performed pca across all index values and derived absolute values for the highest ranked four principal components for each index, since we were interested in overall magnitude, rather than in the direction, positive or negative, along which the components were grouped according to a particular algorithm. Figure 4 illustrates a combined coherence map of the highest loadings for top components across indices. Importantly, the inferior parietal is confirmed as a key area of coherence estimation.

**Figure 4.**
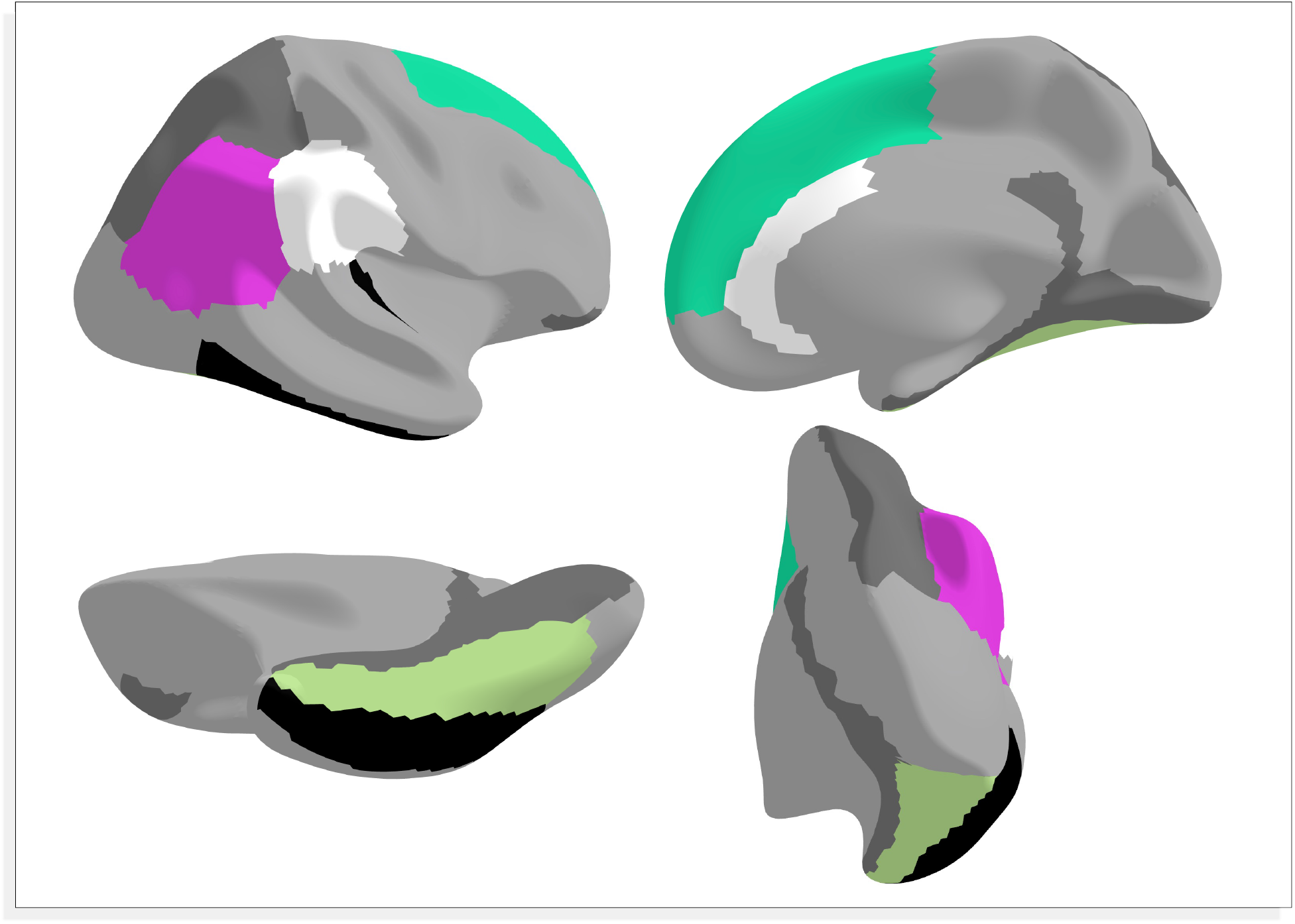
Coherence map based on highest four principal component loadings for each index. Color coding: three shared ROIs across all indices DI, LI, SI, and ZI in magenta (inferior parietal), sea green (superior frontal), and olive green (fusiform); three ROIs shared by LI, SI, and ZI in white (supramarginal, caudal anterior cingulate, rostral anterior cingulate); six ROIs specific to DI in dark grey (entorhinal, lingual, parahippocampal, pars orbitalis, superior parietal, and isthmus cingulate); two ROIs shared among some of the indices LI, SI, and ZI, but not by DI in black (transverse temporal and inferior temporal).

To investigate whether brain coherence is a robust predictor of cognitive performance across diagnosis groups in distinct ways for males and for females, we entered the 25 cognitive test measures into linear models, with the first four principal components of each of the four indices as predictors alongside diagnosis, sex, and their interactions with the four components. We counted 15, 12, 13, and 11 models with significant statistics and meaningful explained variance (*R2* > 0.1), for DI, LI, SI, and ZI respectively, representing 60%, 48%, 52% and 44% out of 25 cognitive measures. In an ANOVA over the 25 models corresponding to as many cognitive measures, DI ranked highest with a mean *R^2^* of 0.191 (*SD* = 0.062), followed by SI with 0.186 (*SD* = 0.058), LI with 0.174 (*SD* = 0.068), and ZI with 0.172 (*SD* = 0.055). However, cognitive score predictions did not differ significantly across indices, *F* (3, 100) = 0.595, *p* = 0.62 – also see Figure 7.

**Figure 5.**
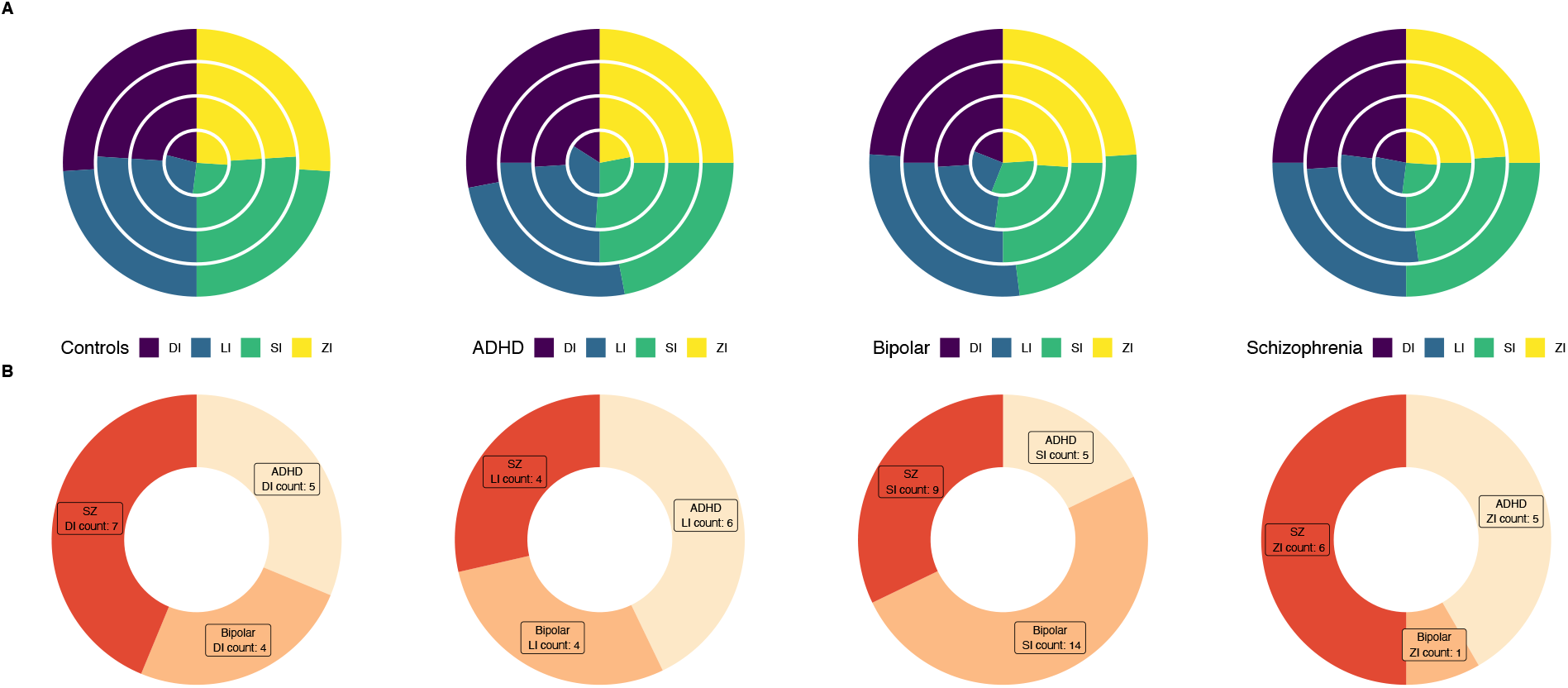
Interactions between DI, LI, SI, and ZI principal components with diagnosis groups as predictors of cognitive measures. Upper quartile magnitudes of effect sizes for the top 20% highest ranked index components (1-to-4 from the center out) entered as predictors in linear models in the is relatively similar across diagnosis groups (ADHD, Bipolar, Schizophrenia, and Controls) and across indices (a). The number of significant effect sizes predicted by interactions between principal components and diagnosis groups compared to female controls is balanced for DI and LI and unbalanced for SI AND ZI (b).

**Figure 6.**
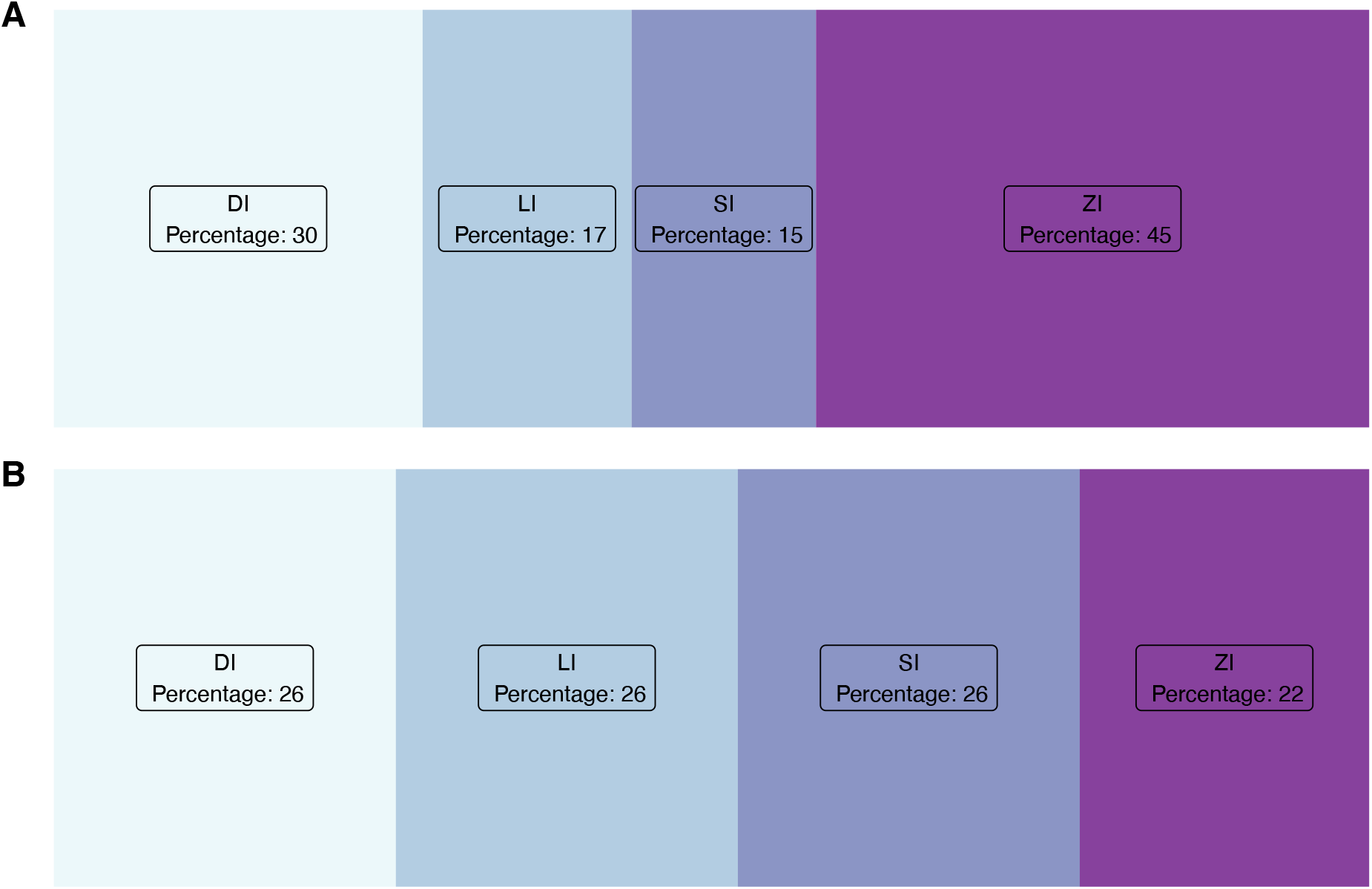
Interactions of DI, LI, SI, and ZI principal components and sex as predictors of cognitive measures. The percentage of significant interactions between sex and indices is highest for ZI components and lowest for LI and SI components, with DI featuring intermediate values (a). The percentage of significant interactions between sex and index components having opposed sign (signaling changes in coherence) compared to female controls out of the total number of significant index-sex interaction. The percentage is halved for ZI, is sensibly increased for LI and SI, and remains stable for DI.

**Figure 7.**
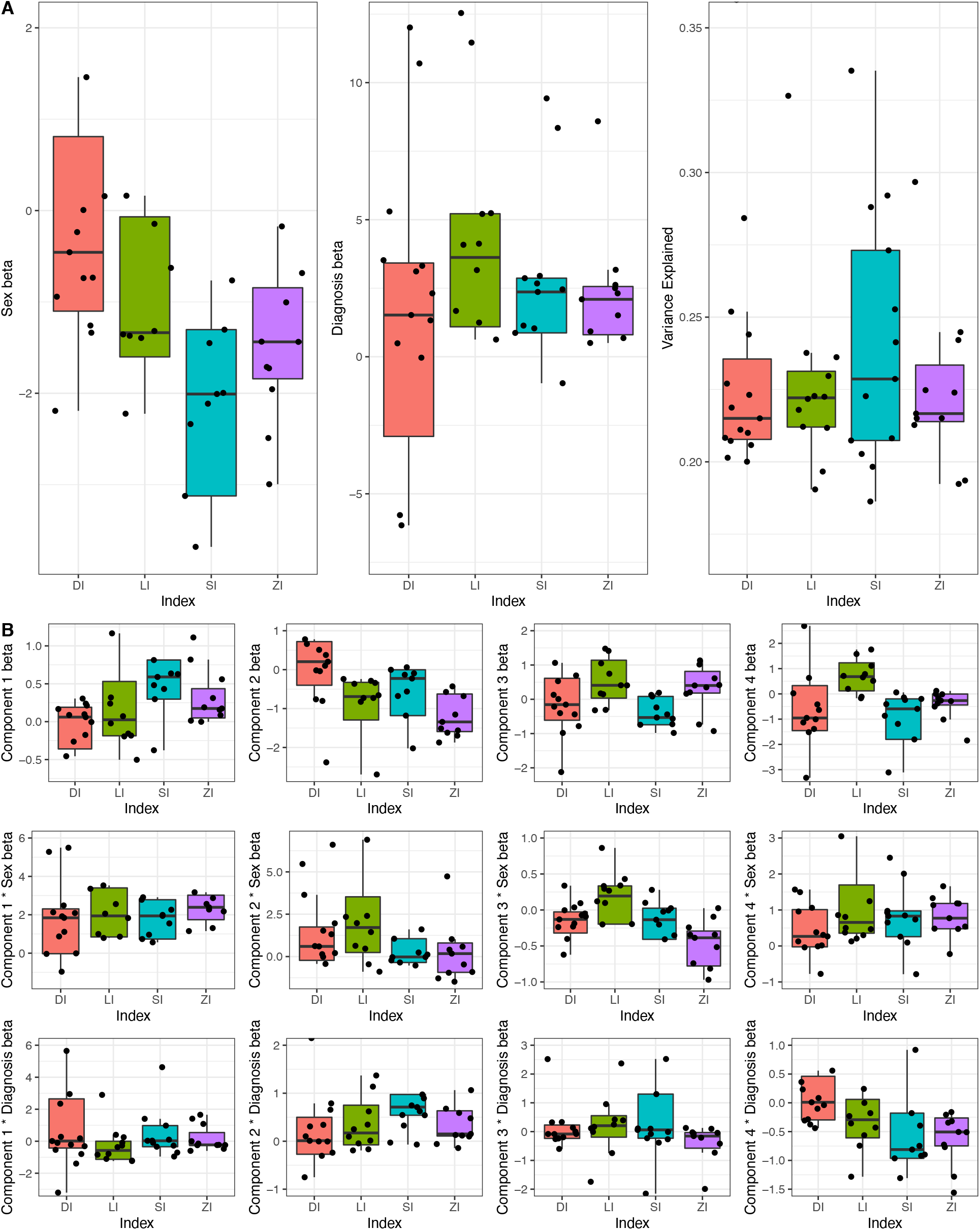
Effects of sex and diagnosis with variance explained for 25 cognitive measures across indices DI, LI, SI, and DI (a) and interaction effects for 25 cognitive measures (b).

Figure 5a compares the magnitude of the highest ranked (upper quartile) model effect sizes for the four principal components of each index. Again, the patterns of effects magnitude across indices are similar for diagnosis groups, with greater differences between DI, LI, SI, and ZI for the first component. Interestingly, effect size magnitudes were similar not only across the four indices, but also across the four components of each index, which underscores the similarity of these four measures of coherence considering that, for each index, there were differences of hundreds of orders of magnitude between the five highest ranked magnitudes and the very last ones among the 25 measures. Further, we can observe greater similarity across indices for controls than for any of the three diagnosis groups. Figure 5b gives an overview of the relative number of significant effect sizes for each of the four indices and for each diagnosis group (ADHD, Bipolar, Schizophrenia) with respect to controls. It becomes clear that the number of significant effect sizes is relatively equally distributed across groups for DI, and to a slightly lesser extent for LI. However, the sensitivity of SI and especially of ZI is much lower. In sum, DI and LI are better biomarkers of psychopathology compared to SI and ZI.

When comparing DI, LI, SI, and ZI performance in terms of number of significant effect sizes resulting from interactions between pca components and sex, ZI scored the highest percentage, followed by DI. LI and SI only scored modest percentages, as seen in Figure 6a. However, when looking at the percentage of significant effect sizes having opposed sign relative to coherence parameters of female controls and out of the total number of significant effect sizes for interactions between index components and sex, ZI percentage is halved. In contrast, we note improved performance for LI and SI, with DI score remaining constant, as seen in Figure 6b. In sum, DI appears to be the most robust biomarker of both sex and psychopathology compared to LI, SI, and ZI based on their performance as predictions of cognitive test measures.

## Discussion

We tested the hypothesis that biomarkers of brain coherence help distinguish male brains from female brains, which in turn serve as models for psychopathological conditions. We developed the distance index as a coherence biomarker over left versus right hemisphere volumes and compared it with three other coherence indices calculated ad hoc: whole-brain absolute laterality index LI, subtraction index SI, z-scored laterality index ZI. We provided support for our hypothesis in several ways, as follows. First, we found that only DI could reliably distinguish between males and females and between Controls and diagnosis groups (ADHD, Bipolar). Since the sample was highly unbalanced for the Schizophrenia group (about four times as many males than females), we did not find a significant difference from Controls. Second, we highlighted a network of cortical regions associated with whole-brain coherence across groups for each index based on score values, which were similar for males and for females, but included a few areas which were swapped between the ADHD group and other groups, as predicted by our male/female brain model hypothesis. Second, we singled out the inferior parietal as a shared coherence area among groups, sexes, and indices, to be explored in future studies. Third, we found significant effect sizes in linear models where index components and their interaction with diagnosis groups predicted cognitive test scores. In other words, we determined that psychopathology impacts brain coherence. However, the four coherence measures were not equally sensitive to diagnosis, as only DI and LI (to a slightly lesser extent) could predict cognitive measures for the three psychopathological conditions with respect to Controls. Fourth, we found coherence to differ between males and females when predicting cognitive test scores, with DI providing the most robust evidence. Specifically, we reported the highest percentage of opposed-sign interactions of DI principal components with the predictor ‘sex’ across the four indices. Indeed, since opposed sign interactions represent the best evidence for a change in brain coherence, DI is a better biomarker of psychopathology.

To summarize, we found that female brains are structurally more coherent than male brains in healthy individuals as well as in patients with bipolar disorder and Schizophrenia based on analyses of DI values. In contrast, based on analyses of index values and shared ROIs, we found a notable exception in the ADHD group, where the reverse was true, with female brains being less coherent than male brains. This reversal confirms previous reports that ADHD is a male-specific illness for having higher prevalence in males, whereas depression has higher prevalence in females (Angold, Costello, and Worthman, 1998). These findings suggest that the brain of individuals suffering from ADHD follows the healthy male model, involving a shift in coherence between the sexes, with the female brain being less coherent than the male brain. This reversal deserves further investigation, as it suggests that the ADHD female brain is structurally an extreme simulation of the male brain, or that changes may concern females rather than males.

Importantly, brain coherence is distinct from brain lateralization, with the former being underspecified for left-right structural differences. Indeed, we defined coherence as a homologous pattern of volumes for left and right anatomical structures, hence each of the 31 brain regions are about as different from each other as the corresponding 31 grey matter volumes of the right hemisphere. The same holds for male and female brains in the three pathological conditions investigated, ADHD, bipolar disorder and schizophrenia, as well as in healthy individuals. Therefore, reduced lateralization does not entail higher coherence, and, conversely, highly lateralized structures are not always less coherent. Indeed, left and right anatomical structures could be equally coherent, as increased coherence of a specific region of interest does not imply a difference in size between the left and the right counterparts. Thus, the volume of one brain hemisphere or for the whole brain may remain unchanged as coherence differs in terms of relative volume of the combined regions of interest.

Finding reliable biomarkers of brain coherence has long been a desideratum for MRI-driven research. Indeed, structural brain-behavior associations (Ismaylova et al., 2018; Kim et al., 2015; Luders et al., 2013; Luders et al., 2012; McEwen et al., 2016) have been described in conjunction with variations in personality, intelligence and even political preferences, but recent attempts to replicate many of these findings have been unsuccessful, thus questioning the existence of a direct link between brain structure and human behavior. For example, increased brain symmetry is known to be a hallmark of psychopathology, which often involves impaired cognitive performance. However, no reliable MRI-based biomarkers were available that could distinguish the brain structure of the mental ill from that of healthy individuals. In particular, Boekel and colleagues (2015) investigated a series of specific previously reported structure-behavior associations but could not find support for the original results in a replication study. Also, Genon and colleagues (2017) reported a number of correlations between cognitive performance and measures of gray matter volume in sub-regions of the dorsal premotor cortex for healthy adults, which nevertheless seemed counterintuitive (i.e., higher performance related to lower gray matter volume).

This ‘replication crisis’ for studies on brain structure and psychological phenotypes (Kharabian et al., 2019) echoes the general replication crisis ramming psychological studies (Button et al., 2013; De Boeck and Jeon, 2018). More to the point, several reasons have been identified for the stalemate of MRI-driven research on biomarkers. They include the relative shortage of longitudinal and developmental studies (Herting et al., 2018, Buimer et al., 2020), the emphasis on unimodal over multimodal imaging, and the constant need for novel analysis techniques. We have argued that targeting the whole brain instead of specific areas of the brain might offer solutions to be investigated in subsequent studies. Computing DI in conjunction with cognitive performance measures to assess brain coherence allows for a more sensitive and encompassing analysis of brain structures specific to individuals within and across the pathological boundary than using other measures such as LI, SI, or ZI. In other words, DI is a promising tool to explain the link between brain structure and behavior based on male versus female structural models, with relevance to both pathological and non-pathological samples.

## Author Contributions

MD contributed the concept and design of the study, data analysis and interpretation, and drafted the manuscript. MK and HB contributed data analysis and interpretation. All authors reviewed the manuscript.

